# Effect of agrochemical exposure on *Schistosoma mansoni* cercariae survival and activity

**DOI:** 10.1101/701060

**Authors:** Devin K. Jones, David D. Davila, Karena H. Nguyen, Jason R. Rohr

**Affiliations:** University of Notre Dame, Department of Biological Sciences, Notre Dame, IN, USA 46556; University of Miami, College of Arts and Sciences, Coral Gables, FL, USA 33146; University of South Florida, Department of Integrative Biology, Tampa, FL, USA 33620

## Abstract

Land conversion and agrochemical use has altered freshwater systems worldwide, introducing chemicals and pathogens (e.g., helminths) that threaten human health. In developing countries where stringent pesticide use and water treatment is limited, understanding how contaminants and pathogens interact is of particular importance. Schistosomiasis, a neglected tropical disease, is caused by the free-swimming cercariae of *Schistosoma mansoni*, a flatworm (trematode) that is transmitted from snails to humans. Schistosomiasis afflicts over 200 million people, reinforces poverty, and has an enormous impact on children. To investigate the effects of pesticide exposure on *S. mansoni*, we exposed cercariae to four insecticides (cypermethrin, deltamethrin, dimethoate, and methamidophos) at five concentrations above estimated environmental concentrations, and recorded survival and activity during a 24-hr time-to-death assay. To identify live, but paralyzed, cercariae from dead cercariae, we used Trypan blue dye, which is only expelled from live cells. We found no effect of cypermethrin, deltamethrin, or dimethoate exposure on the survival and activity of *S. mansoni* cercariae. Surprisingly, methamidophos exposure decreased activity and increased survival of cercariae compared to those in control treatments. This result is likely due to methamidophos causing paralysis of cercariae, which reduced energy consumption lengthening lifespan. Although methamidophos exposure increased survival time, the pesticide-induced paralysis left cercariae functionally dead, which could influence overall disease prevalence and thus human health. Future studies that examine the influence of agrochemicals on waterborne disease prevalence and transmission need to consider both the lethal and sublethal effects of exposure to fully understand the complexity of host-parasite interactions.

**Author Summary:** Previous methods used to investigate the effects of pesticide exposure on free-swimming life stages of trematode pathogens include 1) normal activity, 2) movement following stimuli, or 3) staining dyes. As pesticides commonly target motor function, the use of an individual metric to assign trematode survival might misidentify pesticide-induced paralysis as mortality, therefore underestimating trematode tolerance. In this study, we used activity assays in tandem with Trypan blue staining dye to assess the effects of four pesticides on *Schistosoma mansoni* cercariae. We found that cercariae are highly tolerant to pesticide levels far beyond environmentally relevant concentrations. Surprisingly, exposure to methamidophos increased the survival and decreased the activity of cercariae compared to those in control treatments. Reduced activity was presumably caused by methamidophos-induced paralysis of cercariae. Although we observed increased survival following methamidophos exposure, the pesticide-induced paralysis rendered cercariae functionally dead. Our results highlight the need for future assays examining trematode tolerance to contaminants to employ both activity assays and staining dye to discern cercarial paralysis from mortality. Understanding the effects of pesticide exposure on disease transmission is of vital importance as pesticide use and agricultural activities intensify in developing nations endemic to waterborne pathogens.

## 1. Introduction

The rate of production and use of synthetic chemicals, such as pesticides, has outpaced other human drivers of global environmental change (1), which has resulted in the contamination of ecosystems worldwide (2–4). Exposure to pesticides can cause direct, lethal effects on sensitive species or sublethal effects on an organism’s behavior, physiology, and/or morphology (5–10). Organophosphate and pyrethroid insecticides, for example, target important esterases and nerve cell gates leading to ionic imbalances and uncontrollable convulsions and tremors before paralysis and eventual death (11, 12). Given that pesticide production and trade is estimated to increase drastically by 2050 (13, 14), there is a growing need to understand how agrochemical contamination impacts human and wildlife health.

Freshwater ecosystems, which are vital for global economies, societal well-being, and maintaining human health (15–17), are threatened by agricultural activities and agrochemical use (4, 18, 19). For instance, over 40% of the global land area is at risk of producing insecticide runoff to lotic systems (19). This same agricultural runoff can also carry microorganisms including bacterial, viral, fungal, and helminth pathogens that cause waterborne diseases, further jeopardizing human health (20, 21). In developing countries, such as those in sub-Saharan Africa, increased land conversion for agriculture (13, 22) combined with water management and development for irrigation (23, 24) has led to an increase in human exposure to agrochemicals and waterborne pathogens (25–27). Understanding the effects of agrochemical contamination on waterborne diseases is vital as the density of human settlements near managed water systems and the demand for agricultural output are increasing (14, 23, 28).

Schistosomiasis is one example of a waterborne, neglected tropical disease that is affected by agriculture. Schistosomiasis afflicts over 200 million people, of which over 90% reside in sub-Saharan Africa (29, 30), and is caused by parasitic *Schistosoma* trematodes. Free-swimming *Schistosoma* cercariae are released from intermediate snail hosts and penetrate the skin of definitive human hosts while in the water. *Schistosoma* eggs, produced by matured worms, leave the human host via feces or urine, and hatch in aquatic environments where the next free-living stage, miracidia, penetrate the snail intermediate host to complete the life cycle. Cercariae and miracidia are short-lived organisms, generally only having enough reserves to live for approximately 24 hours (31). In an outdoor mesocosm experiment, populations of intermediate snail hosts were shown to increase following bottom-up and top-down indirect effects caused by herbicide and insecticide exposure, respectively (32). Thus, human infection risk was predicted to increase following pesticide contamination of aquatic systems. As pesticide runoff potential is high in developing African nations where schistosomiasis is prevalent (19, 29, 33), investigating the direct effects pesticide exposure has on the aquatic life stages of *Schistosoma* is of great importance.

*Schistosoma* miracidia and cercariae might be sensitive to environmental contaminants given their occurrence in freshwater environments during transition between intermediate and definitive hosts (34). Surprisingly, previous studies have not reported significant lethal effects of pesticide exposure on either life stage of *Schistosoma* (32). In contrast, similar trematode species found in North American snails are known to be sensitive as cercariae to the commonly applied herbicides atrazine (35, 36) and glyphosate (37), as well as organophosphate, pyrethroid, and neonicotinoid insecticides (38). Interestingly, the method of assigning cercarial survival differed among these studies. Survival can be assessed with 1) general swimming or climbing movement (36), 2) movement following stimuli (35, 38–40), or 3) Trypan blue staining (32). Complicating the use of either activity or dyes to examine effects of pesticide exposure is that many insecticides cause paralysis; thus, paralyzed cercariae cannot respond to stimuli but can excrete Trypan blue, which is absorbed by dead cells and excreted by living cells (41, 42). If exposure to insecticides that target the nervous system reduce activity or cause paralysis, this should reduce energy consumption of short-lived, free-living, aquatic organisms, such as cercariae, potentially extending their lifespan despite the fact that they are not functional and therefore not infective. This could explain some of the variability in cercarial responses to insecticides.

To assess this hypothesis, we exposed *Schistosoma* cercariae to five different concentrations of four insecticides from two different pesticide classes (organophosphate, pyrethroid) using a 24-hr time-to-death (TTD) assay. We focused on organophosphate and pyrethroid insecticides because they are nerve agents that can disrupt muscle activity and thus movement. To distinguish among active cercariae, paralyzed cercariae, and dead cercariae, we simultaneously employed activity assays and Trypan blue staining (S1 Fig). We predicted that cercarial exposure to the insecticides would cause paralysis, thus reducing activity and extending survival compared to cercariae not exposed to insecticides that were actively swimming. We also predicted similar effects on cercarial activity regardless of organophosphate or pyrethroid exposure given the need for both acetylcholine esterase (organophosphate target) and voltage-gated ion channel (pyrethroid target) function in *Schistosoma* cercarial movement (43, 44).

## 2. Methods & Materials

### 2.1 Pesticide background

We chose four insecticides commonly used and detected in sub-Saharan African countries (27, 45, 46). Dimethoate (CAS 60-51-5) and methamidophos (CAS 10265-92-6) are broad-spectrum organophosphate insecticides that are acetylcholinesterase (AChE) inhibitors used to protect crops such as grapes, tobacco, and potatoes. Deltamethrin (CAS 52918-63-5) and cypermethrin (CAS 52315-07-8) are Type-II pyrethroid insecticides that mimic natural pyrethrins by interfering with sodium ion channels of nerve cells. Deltamethrin and cypermethrin are applied to numerous agricultural crops, such as corn, cotton, and rice, and play a vital role in integrated pest management strategies to reduce vector populations. Although estimated use records of each pesticide are scarce for African countries, their use is listed by the Pesticide Action Network Africa (http://www.pan-afrique.org/departen.php). Moreover, previous research has reported human exposure to each pesticide, and has found residues of each in the air, water, soil, and produce of African countries (46–49).

### 2.2 Study organisms

*Schistosoma mansoni*, the trematode species responsible for intestinal schistosomiasis, is found within aquatic environments in South America, the Caribbean, and Africa. In Sub-Saharan Africa, the increased management of waterways and construction of dams and irrigation channels has contributed to the increased abundance and distribution of intermediate snail hosts, increasing human risk of exposure to schistosomiasis (23, 24, 33).

We obtained 50 infected *Biomphalaria glabrata* snails on September 11, 2018 that were exposed to *S. mansoni* (NMRI strain) on September 5, 2018 from the NIAID Schistosomiasis Resource Center of the Biomedical Research Institute (Rockville, MD). Snails were held individually in 200-mL containers filled with 200 mL HHCOMBO water (Baer et al 1999) and fed *ad libitum* a ration of ground fish flakes (Tetramin®; Blacksburg, VA, USA) and spirulina (NOW FOODS^®^; Bloomingdale, IL, USA) suspended in agar (Fisher BioReagents^®^; Fair Lawn, NJ, USA). Snails were held under laboratory conditions (25.5°C, 12:12 light:dark), and full water exchanges were conducted biweekly.

### 2.3 Time-to-death assay design

We examined the effects of the four pesticides on the survival and behavior of free-swimming cercariae of *S. mansoni* using a 24-hr time-to-death (TTD) assay. We conducted the TTD assay using 24-well tissue culture plates (Falcon® # 353047; Corning Incorporated, Corning, NY, USA). We tested five concentrations of each insecticide for a total of 20 pesticide treatments. To these treatments, we added a water control and an ethanol vehicle control. A vehicle control was included in the experimental design because pyrethroid insecticides are insoluble in water. We included two replicates of each control treatment and one replicate of each pesticide treatment on each 24-well plate and used five plates for a total of 120 wells.

To obtain *Schistosoma* cercariae, eight infected snails were transferred to 50-mL glass beakers filled with 15 mL of oxygenated HHCOMBO water and were held under direct artificial light for 1.5 hr. Snails were then returned to their respective husbandry containers, the 15-mL HHCOMBO solutions containing shed cercariae were homogenized, and we dispensed 250 µL cercariae slurry to each well. On average, this resulted in 5.05 ± 0.43 (mean ± 1 SE) cercariae per well.

We created our pesticide treatments by first making stock solutions of each chemical. Organophosphate insecticides were dissolved directly in HHCOMBO water (5 mg/mL), whereas pyrethroid insecticides were dissolved using ethanol (0.05 mg a.i./mL). We then added an aliquot of each stock solution to 10 mL of HHCOMBO water to create an intermediate solution for each targeted concentration (20 intermediate solutions). Prior to addition of stock solutions, we removed the same volume of HHCOMBO water from the 10-mL intermediate vial that we would be adding to correct for total volume. We added 100 µL of each intermediate solution to their respective wells to obtain the nominal concentrations of 10, 30, 50, 70, and 100 µg/L for pyrethroids and 100, 200, 300, 400, 500 mg/L for organophosphates. Although the chosen nominal pesticide concentrations fall above expected environmental concentrations (Table 1), they were selected following a series of pilot studies with the aim of causing increased cercariae mortality. We attempted to narrow our range of concentrations for each class using pilot studies that employed 0.1, 0.5, 1.0, 2.0, and 10.0 µg active ingredient/L for pyrethroids and 5, 10, 35, 75, and 100 mg active ingredient/L for organophosphates. However, we did not observe significant death at these lower concentrations when compared to the water control. Thus, concentrations were increased for both pesticide classes in the final experiment. The ethanol vehicle control was created by adding 101 µL of ethanol (95%) to 9.899 mL of HHCOMBO water to match the ethanol concentration in the highest volume of pyrethroid stock solution being transferred to the intermediate solution. To create our water controls, we instead added 100 µL of HHCOMBO water to each respective well. We then added 15 µL Corning™ Trypan blue dye (CAT MT25900CI, Fisher Scientific) to each well for cercarial staining. We conducted a 24-hr TTD assay to compare survival of cercariae exposed to Trypan blue stain to that of cercariae in water controls and found no effect of staining on survival (*p* = 0.44). Lastly, we added 135 µL of HHCOMBO water to each well to bring the total volume to 500 µL.

**Table 1.** Peak estimated environmental concentrations (EEC) for each insecticide. We used the USEPA Pesticide in Water Calculator (PWC; version 1.52) to calculate EEC for pond and reservoir surface waters. Following previously described methods (63), we extracted pesticide parameters from the University of Hertfordshire’s Pesticide Properties DataBase (PPDB; https://sitem.herts.ac.uk/aeru/ppdb/en/), the Pesticide Action Network (PAN) Pesticide Database (http://www.pesticideinfo.org/), and the Hazardous Substances Data Bank (HSDB; https://toxnet.nlm.nih.gov/cgi-bin/sis/htmlgen?HSDB). We then selected the maximum EEC value generated by the PWC calculator.

| Pesticides | Crop | Water body | Peak EEC<br>(ppb; ug/L) |
| --- | --- | --- | --- |
| Methamidophos | Potato | Pond | 6.05 |
|  |  | Reservoir | 13.9 |
| Dimethoate | Corn | Pond | 8.17 |
|  |  | Reservoir | 19.2 |
| Cypermethrin | Cotton | Pond | 0.859 |
|  |  | Reservoir | 2.03 |
| Deltamethrin | Corn | Pond | 0.0036 |
|  |  | Reservoir | 0.0086 |

We assessed survival and activity of cercariae during the 24-hr toxicity test. We counted the number of unstained (alive), stained (dead), and active cercariae every two hr for the first 12 hr, and then every six hr for the second 12 hr. Activity was recorded if cercariae were actively swimming, crawling, or moving in the water column. We did not conduct water exchanges or renew pesticide concentrations during the 24-hr exposure period. After 24 hr, we added 20 µL Lugol’s iodine solution to each well to euthanize and stain surviving cercariae. Lugol’s iodine solution was used to determine the total number of cercariae per well as we used a standardized volume of shed cercariae in favor of separating individuals to reduce handling time of cercariae.

### 2.5 Statistical analysis

To examine the direct toxic effects of the four insecticides on *S. mansoni* cercariae, we analyzed cercarial survival over time using Cox’s proportional hazard models (50). We first conducted an analysis comparing survival of cercariae exposed to the ethanol vehicle control and the water control to assess any effect of the vehicle. We did not find any difference between the two treatments (*p* = 0.94; S1 Table). We thus pooled the ethanol vehicle and water controls for all subsequent survival analyses. We then conducted four independent survival analyses, one for each insecticide, to examine the effect of concentration (continuous variable) on cercarial survival. The pooled control treatment served as a 0.0 µg/L concentration in each model. Following a significant effect of concentration, we then compared the survival of cercariae in each insecticide concentration (categorical variable) to the survival of cercariae in the pooled control treatment. We included ‘experimental well’ as a random effect in each model. Cox’s proportional hazards model were employed using RStudio Version 1.1.453 (51) and the *survival* and *coxme* packages. Additionally, we used the *drc* package in RStudio to estimate the effective dose (ED10, ED50, and ED90) for pesticides that induced significant concentration effects on cercarial survival. We first used the *drm* function to examine the effect of log_10_-transformed methamidophos concentration (+ 1) on the occurrence of cercarial death, and then back-calculated estimated effective doses (10^X^ −1).

To test whether cercarial activity over time was affected by insecticide exposure, we employed generalized linear mixed-effects (GLME) models. We first examined if activity over time differed between cercariae exposed to the water and ethanol vehicle controls. We examined if the interactive effects of control treatment and time (independent variables) influenced the activity of cercariae, represented by the binomial response of the number of active and inactive cercariae within each experimental well. We found no difference in the activity of cercariae in the two control treatments (χ^2^ (1) = 1.97, *p* = 0.161), so we pooled the water and ethanol vehicle controls. For each insecticide, we then investigated the interactive effects of concentration (continuous) and time (independent variables) on cercarial activity. If we observed a significant effect of concentration, we then conducted a subsequent model that investigated the main and interactive effects of pesticide concentration (categorical) and time on the activity of cercariae and conducted Tukey’s post-hoc pairwise comparisons. We included ‘experimental well’ as a random effect term within each model. Model analyses were conducted using RStudio and the *car*, *lme4*, and *multcomp* packages.

## 3. Results

### 3.1 Time-to-death assays

Cox’s proportional hazard models revealed that there was no effect of concentration on the survival of cercariae exposed to cypermethrin, deltamethrin, or dimethoate (*p* ≥ 0.64; S1 Table, S1 Dataset). In contrast, we did find a significant effect of methamidophos concentration on cercarial survival (*b* = −0.003, *p* = 0.002). Exposure to 100, 200, 300, and 400 mg/L methamidophos increased cercarial survival relative to the pooled controls (*p* ≤ 0.031; Fig 1, S1 Table). Survival of cercariae exposed to 500 mg/L methamidophos did not differ from survival of cercariae in the pooled controls (*p* = 0.12). After 24-hr of exposure, survival of cercariae in the pooled control was 41.3% compared to >73% for cercariae exposed to any methamidophos concentration.

**Fig 1.**
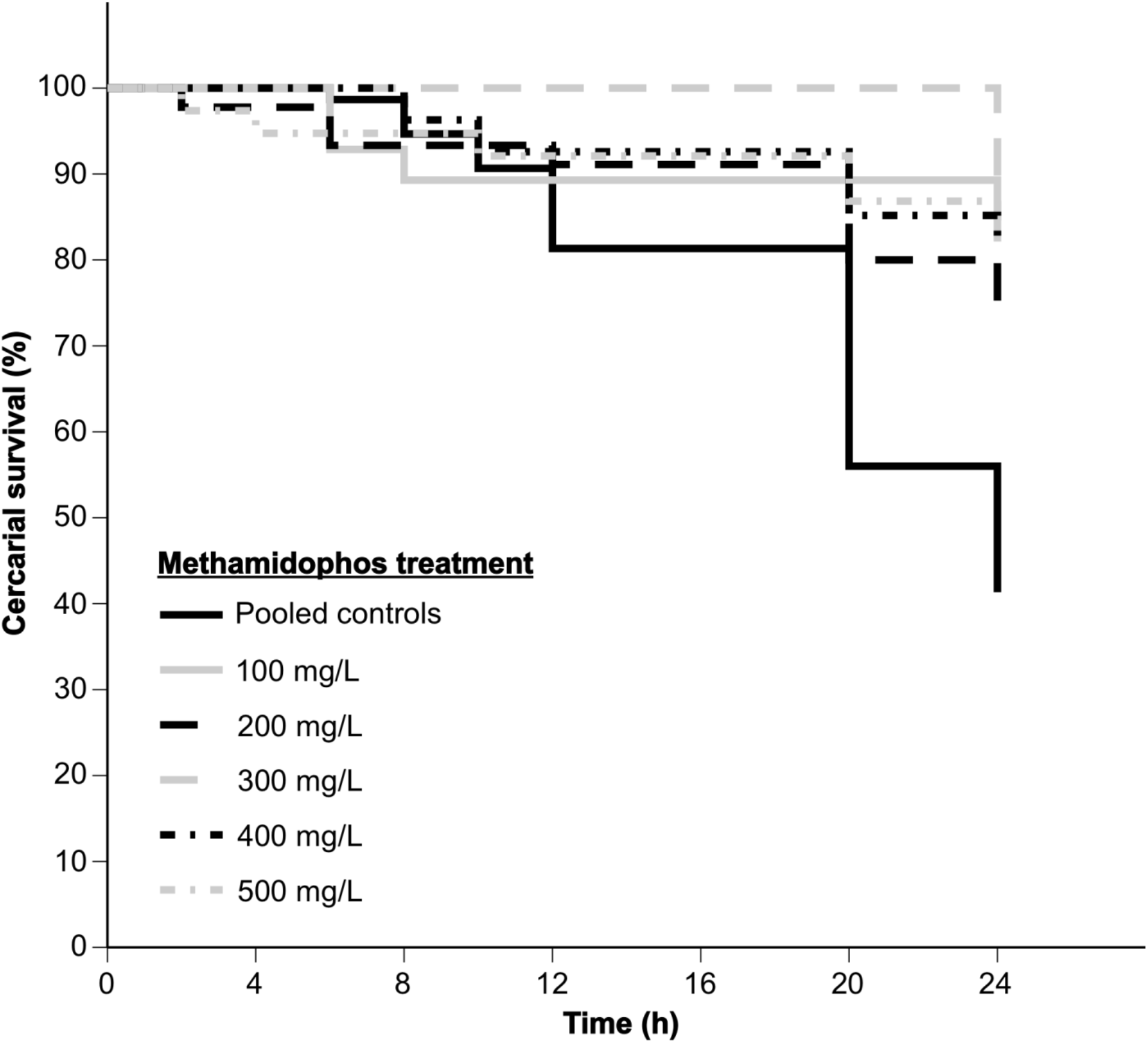
Survival of *S. mansoni* cercariae following exposure to one of six methamidophos concentrations. Cercariae were exposed to 0, 100, 200, 300, 400, or 500 mg/L methamidophos for 24 hr using a time-to-death assay. The pooled control treatment represents the combined survival of cercariae in the water and vehicle control treatments (*p* = 0.94).

To examine the toxicity of methamidophos to *S. mansoni* cercariae, we calculated the 24-hr effective dose. The slope (b; *p* = 0.4517) and LD50 (e; *p* = 0.4380) parameter estimates from the two-parameter log-logistic model with fixed lower and upper limits were not different from zero. The estimated 24-hr 10, 50, and 90% effective doses (± SE) for methamidophos were 0.61 (± 3.68), 7.06 (± 13.75), and 9770.92 mg/L (± 628.13), respectively.

### 3.2 Activity assay

To examine the influence of insecticide exposure on cercarial activity, we used generalized linear mixed-effects models. While we did not find an effect of concentration (*p* ≥ 0.149) or a concentration-by-time interaction (*p* ≥ 0.366) for dimethoate, cypermethrin, or deltamethrin, activity declined with time for all three insecticides (*p* < 0.001). For methamidophos, concentration (χ^2^ (1) = 37.48, *p* < 0.001) and time (χ^2^ (1) = 46.40, *p* < 0.001), but not their interaction (χ^2^ (1) = 0.534, *p* = 0.4649), influenced cercarial activity (Fig 2, S2 Dataset). Post-hoc multiple comparison tests (Tukey) revealed that mean activity of cercariae exposed to 300, 400, and 500 mg/L methamidophos was lower than the activity in the pooled controls (*p* ≤ 0.047; Fig 2A). Activity of cercariae exposed to 100 and 200 mg/L methamidophos did not differ from that in the pooled controls (*p* ≥ 0.076). Excluding the 0 mg/L control, activity of cercariae did not differ among methamidophos concentrations (*p* ≥ 0.071). Mean cercarial activity declined over time (Fig 2B).

**Fig 2.**
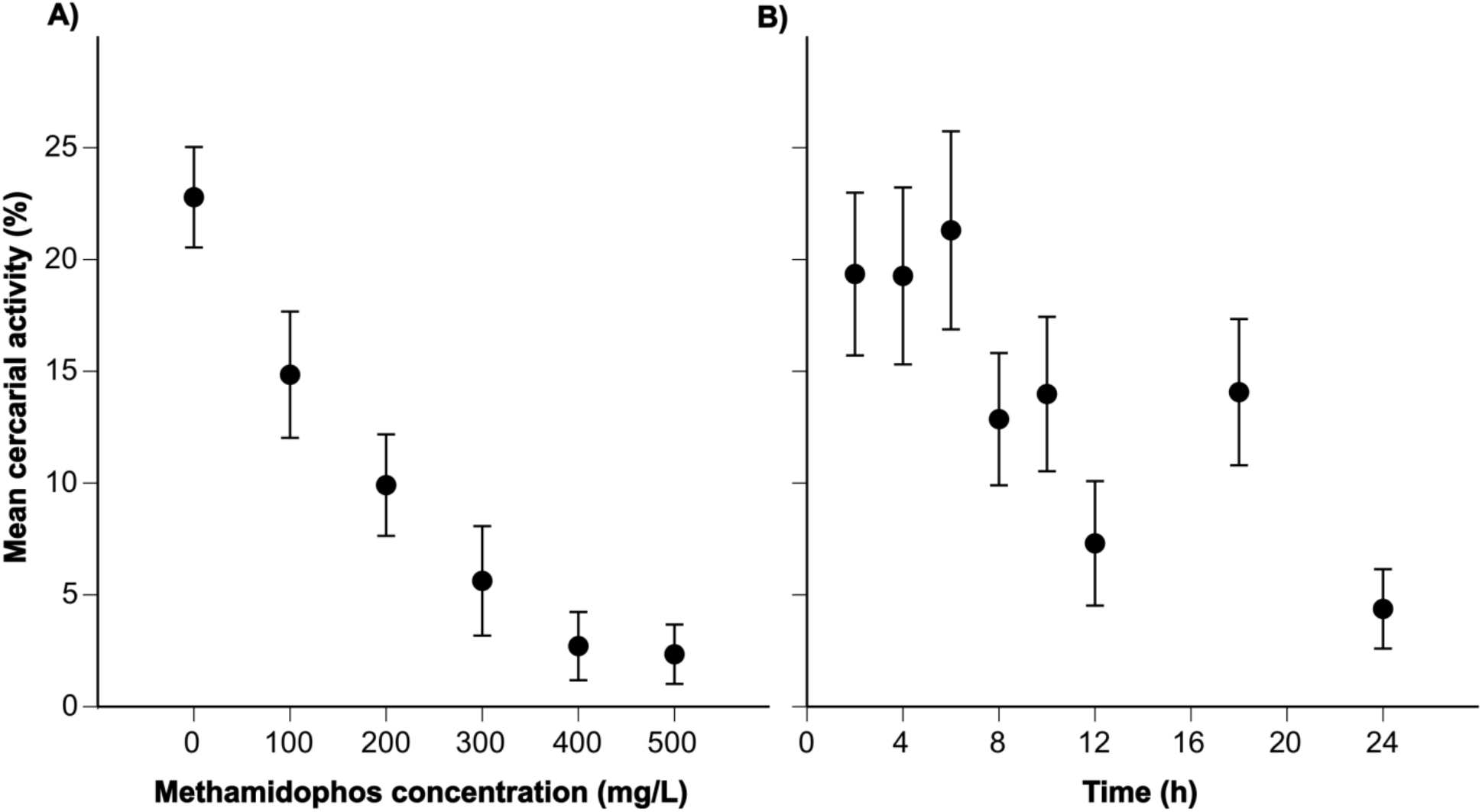
Mean cercarial activity (%) of *S. mansoni* following exposure to one of six methamidophos concentrations. We recorded the number of cercariae active over 24 hrs following exposure to 0, 100, 200, 300, 400, or 500 mg/L methamidophos. We calculated cercarial activity (%) by dividing the number of trematodes observed moving by the total number of individuals in each respective well. We observed the main effect of methamidophos concentration (A) and time (B) on cercarial activity. Data points represents overall treatment mean values ± 1 SE.

## 4. Discussion

In the current study, we sought to understand how free-swimming *Schistosoma* cercariae respond to four insecticides from two chemical classes commonly used in agricultural practices in developing regions endemic to schistosomiasis. We found no significant effect of exposure to cypermethrin, deltamethrin, and dimethoate on the survival and activity of *S. mansoni* cercariae over 24 hr. Surprisingly, exposure to methamidophos resulted in increased cercarial survival compared to the pooled control treatments. Moreover, the use of activity assays in combination with Trypan blue staining allowed us to observe that this increased survival appeared to be caused by the reduced activity of cercariae exposed to methamidophos, which extended their life.

Understanding the influence of common-use pesticides on waterborne diseases is vital to protecting human health in developing regions. Our results suggest that cercariae are highly tolerant to the direct toxic effects of cypermethrin, deltamethrin, dimethoate, and methamidophos contamination. Given that the concentrations of insecticides reported in samples from developing regions all fall below concentrations used in the current study (46, 47, 52–54), it is unlikely that *Schistosoma* cercariae suffer direct mortality from pesticide exposure. Previous research has also reported no influence of chlorpyrifos (organophosphate) or atrazine (triazine) exposure at environmentally relevant concentrations on *S. mansoni* survival over 12 hr (32). While the indirect effects of agrochemicals have been shown to potentially propagate schistosomiasis (32), the results of this study suggest that, due to substantial cercarial tolerance, direct toxicity of pesticides is not an apparent counteractive or mitigating factor for disease risk. As it is unlikely that cercariae in surface waters of natural systems are exposed to only a single chemical compound at a time, future research should investigate the effects of pesticide mixtures on cercarial longevity and infectivity (8). Moreover, investigation into the direct toxic effects of other pesticide classes, such as organochlorines, will be useful as older, more toxic pesticides are still used in developing regions due to availability and low cost (55).

Exposure to pesticides can also cause sublethal changes in behavior and physiology that can alter metabolic processes and energy use. We observed decreased activity of *S. mansoni* cercariae exposed to methamidophos, which was likely caused by full or partial paralysis. The life span (24-48 hr) of *S. mansoni* cercariae is dependent on finite glycogen and fat reserves, and we hypothesize that the paralyzed cercariae might have prolonged longevity because of lower rates of energy consumption (56, 57). Indeed, we observed reduced cercarial activity among individuals exposed to methamidophos relative to the activity of cercariae in all other treatments. Other pesticides and naturally occurring chemicals have also been reported to reduce mobility of nematodes and trematodes, including *S. mansoni* (36, 58, 59). We believe that the exposure to methamidophos caused a true paralytic effect through acetylcholinesterase inhibition in affected individuals, as past research has shown certain cholinergic agents exert an inhibitory effect on muscular activity of *S. mansoni* and other parasites (60, 61).

Although methamidophos-exposed cercariae lived longer than cercariae in the pooled controls, they were “functionally dead” as their immobility prevents them from searching for and infecting definitive hosts (59). Therefore, the methamidophos-induced paralysis could reduce disease transmission and negative impacts on humans. Infection assays in the future should seek to confirm whether cercarial paralysis observed in this study provides protection to human hosts (59). Lastly, the methamidophos-induced paralysis may be strain-, species-, and life stage-specific, providing many questions for future researchers including the comparison of tolerance between lab-reared and field-collected *Schistosoma* cercariae, and if tolerance varies among free-swimming cercariae or miracidia and encysted individuals.

The combined use of Trypan blue staining and activity assays employed in the current study not only allowed us to discriminate whether cercariae were active versus inactive, but also whether they were paralyzed versus truly dead. This could have implications for the interpretation of previous toxicological assays of free-swimming trematode life stages. For instance, previous research using activity to assign mortality to paralyzed cercariae might underestimate the actual tolerance of trematodes to pesticide concentrations (35). Moreover, if the effects of pesticide exposure are short-lived and reversible, either through host metabolism, environmental breakdown, or clearance by flowing water, we may overestimate the toxicological effects of pesticides on disease transmission (32). In natural systems, it is possible that the continuous daily release of thousands of cercariae by infected snail hosts (62) will minimize the influence of pesticide-induced cercarial paralysis on disease dynamics. Alternatively, if pesticide exposure overlaps with peak hours of cercarial shedding, pesticide-induced paralysis might reduce overall infection risk. Future research that examines the complex relationship among infected snail hosts, timing of pesticide exposure, and environmental conditions (e.g., flow, temperature) will reveal how each contributes to the prevalence and transmission dynamics of schistosomiasis (*sensu* (32)).

Freshwater systems are increasingly threatened by numerous anthropogenic activities (3, 17, 23). Understanding the effects of contaminants on waterborne disease is of utmost importance in developing nations where water scarcity and increased agricultural activity might threaten human health (14). Given the increased risk of agricultural runoff in these nations (19), it will be important for future studies to investigate the acute lethal and chronic sublethal effects of contaminants on waterborne pathogens to more thoroughly understand their effects on disease dynamics.

## 5. Acknowledgements

The authors would like to thank Caleb Castro, Karlee Gionet, and Amber Carey for their assistance with snail husbandry and experimental work, and Dr. Samantha Rumschlag for her assistance with statistical analyses and web tool calculations. The University of South Florida’s Leadership Alliance Fellowship awarded to DD enabled this research. *B. glabrata* snails provided by the NIAID Schistosomiasis Resource Center of the Biomedical Research Institute (Rockville, MD) through NIH-NIAID Contract HHSN272201700014I for distribution through BEI Resources. Funds were provided by grants to J.R.R. from the National Science Foundation (EF-1241889, DEB-1518681, IOS-1754868) and the National Institutes of Health (R01TW010286-01).

## Author contributions

JRR, DDD, and DKJ conceived the experimental ideas; DDD, DKJ, and KHN designed the methodology; DDD, DKJ, and KHN collected the data; JRR and DKJ analyzed the data; DKJ and DDD led the writing on the manuscript. All authors contributed critically to the drafts and gave final approval for publication.

## Supporting Information

**S1 Table.** Results of Cox’s proportional hazard models examining survival of *S. mansoni* cercariae following exposure to five concentrations of four commonly used insecticides in Africa. We first compared survival in control treatments before using a pooled control survival in each subsequent analyses. We then determined the effect of concentration of each pesticide. For pesticides with a significant concentration effect, we then compared cercarial survival in each concentration to that in the pooled control treatments. Each model included experimental well as a random effect. Hazard regression coefficient (*b*) and *p-*values are reported.

**Control treatment survival**
|  | *b* | p-value |
| --- | --- | --- |
| Water control<sup>a</sup> | - | - |
| Ethanol vehicle control | -0.023 | 0.94 |

<sup>a</sup>Each treatment was compared to the first in each model.

**Effect of concentration\* on survival**
|  | *b* | p-value |
| --- | --- | --- |
| Methamidophos conc. | -0.003 | **0.002** |
| Dimethoate conc. | < 0.001 | 0.8 |
| Cypermethrin conc. | < 0.001 | 0.98 |
| Deltamethrin conc. | 0.001 | 0.64 |

\* Pooled control survival was used as a 0.0 mg/L pesticide concentration in each model

**Effect of concentration within pesticide treatment on survival**
|  | *b* | p-value |
| --- | --- | --- |
| *Methamidophos* |  |  |
| Pooled control<sup>a</sup> | - | - |
| 100 mg/L | -1.404 | **0.013** |
| 200 mg/L | -1.063 | **0.031** |
| 300 mg/L | -1.722 | **0.028** |
| 400 mg/L | -1.434 | **0.012** |
| 500 mg/L | -1.013 | 0.120 |

<sup>a</sup>Each pesticide concentration was compared to the first treatment in the model.

**S1 Fig.**
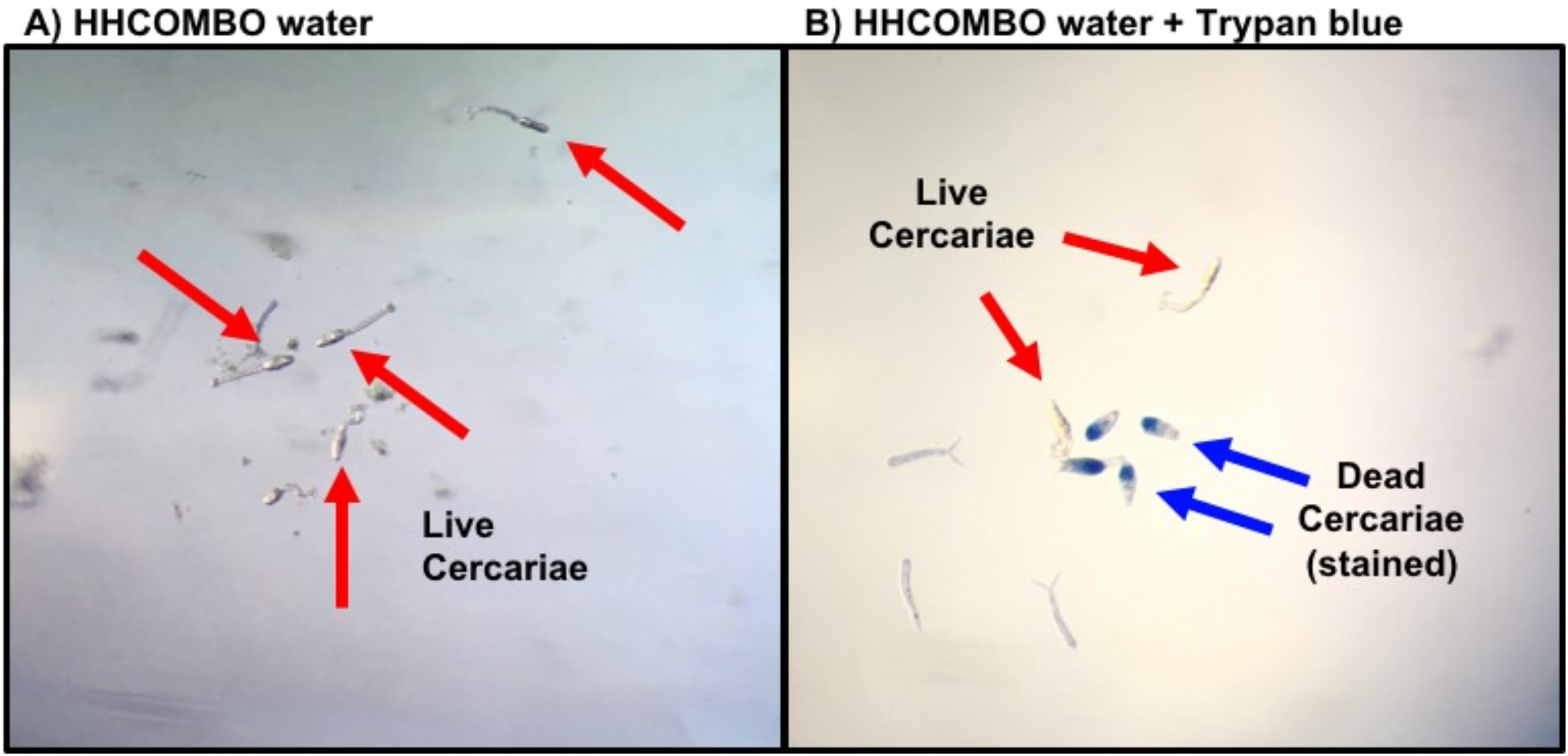
Distinguishing dead cercariae from paralyzed cercariae. When using HHCOMBO water alone (A), observers are unable to identify dead from paralyzed cercariae. We used Trypan blue staining dye (B), which stains dead tissue but is excreted by live cells, to ascertain true death from pesticide-induced paralysis.

### Supporting Datasets

**S1 Dataset. Survival data for the 24-hr time-to-death assay.**

We exposed *Schistosoma mansoni* cercariae to five concentrations of four insecticides and assessed time-to-death over a 24-hr period. The “S1 Dataset_Data_Survival_JonesEtAl.csv” includes survival data for all cercariae in the corresponding study. The values for the column ‘*Event*’ correspond to mortality events; 0 = surviving, 1 = mortality event.

**S2 Dataset. Activity data for the 24-hr time-to-death assay.**

We exposed *Schistosoma mansoni* cercariae to five concentrations of four insecticides and assessed activity over a 24-hr period. The “S2 Dataset_Data_Activity_JonesEtAl.csv” includes cercarial activity data for the corresponding study.

